# A Novel Method and Simple On-line Tool for Maximum Likelihood Calibration of Immunoblots and other Measurements that are Quantified in Batches

**DOI:** 10.1101/026005

**Authors:** Steven S. Andrews, Suzannah Rutherford

## Abstract

Experimental measurements require calibration to transform measured signals into physically meaningful values. The conventional approach has two steps: the experimenter deduces a conversion function using measurements on standards and then calibrates (or normalizes) measurements on unknown samples with this function. The deduction of the conversion function from only the standard measurements causes the results to be quite sensitive to experimental noise. It also implies that any data collected without reliable standards must be discarded. Here we show that a new “1-step calibration method” reduces these problems for the common situation in which samples are measured in batches, where a batch could be an immunoblot (Western blot), an enzyme-linked immunosorbent assay (ELISA), a sequence of spectra, or a microarray, provided that some sample measurements are replicated across multiple batches. The 1-step method computes all calibration results iteratively from all measurements. It returns the most probable values for the sample compositions under the assumptions of a statistical model, making them the maximum likelihood predictors. It is less sensitive to measurement error on standards and enables use of some batches that do not include standards. In direct comparison of both real and simulated immunoblot data, the 1-step method consistently exhibited smaller errors than the conventional “2-step” method. These results suggest that the 1-step method is likely to be most useful for cases where experimenters want to analyze existing data that are missing some standard measurements and where experimenters want to extract the best results possible from their data. Simple open source software for both methods is available for download or on-line use.

**Author Summary:** Most quantitative measurements do not return the physical quantities that are of interest, but some instrument-specific response value instead. These measurements are then converted to physical quantities through a conversion function, which the experimenter deduces from instrument responses for one or more standard samples of known composition. This is called calibration or normalization. For example, we recently performed quantitative immunoblotting on a large number of samples, each replicated on multiple blots, and then calibrated the measurements to yield protein concentrations relative to those in a standard sample. We found that the conventional calibration approach of treating the samples in each blot independently of the samples in other blots produced inaccurate results because this approach is completely dependent on the standard measurements, which were sometimes missing or erroneous in our data. Thus, we developed a new calibration approach in which we fit a statistical model to the entire data set simultaneously. This method, which applies to a very wide range of calibration problems, was substantially more accurate during validation tests and can be shown to return the most accurate results possible within the assumptions of the model. It is particularly useful when some standard measurements are missing from data sets or when experimenters want the best possible results.

## Introduction

Nearly every quantitative experiment requires calibration – the mathematical conversion of raw measurements into physically meaningful values. For example, calibration of immunoblot (Western blot) data converts the intensities of protein bands that are detectable on a blot into the concentrations of proteins that were present in the original samples. Although many scientists take calibration for granted, we show here that conventional approaches are not particularly accurate, causing them to lose some of the information that is carried by valuable measurement data. We present a novel approach that exploits all available information in the data and returns the most accurate results possible within the constraints of a statistical model.

The classical solution to the linear calibration problem [1-4] is a two step process: first, during the calibration step, measurements on known samples, “standards,” are used to deduce a conversion function. Then, during the prediction step, the conversion function is used to convert measurements on unknown samples to physical quantities.

For example, suppose a chemist uses an instrument whose response is linear in the amount of protein, chemical, or other analyte in a sample. This means that an instrument measurement, *y*, is related to the amount of analyte, *x*, according to the response function

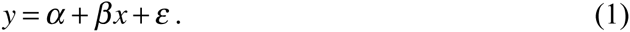

The *α* and *β* parameters are instrument-specific sensitivity coefficients and *ε* represents random measurement noise. In the calibration step, the chemist estimates the *α* and *β* sensitivity coefficients, yielding *a* and *b* respectively, by measuring several standards with known compositions and fitting the resulting data with eq. 1 using linear regression. Substituting the regression results into eq. 1 and solving for *x* yields the conversion function

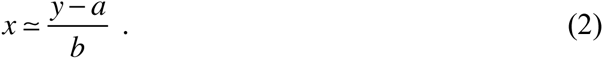

In the prediction step, the chemist measures samples of unknown composition on the same instrument and inserts the measurements into eq. 2. This yields the sample analyte amounts.

In this example, note that errors in the standard measurements lead directly to errors in the sensitivity coefficient estimates. From there, they lead to errors in the computed analyte amounts. For this reason, it is good practice to measure standards repeatedly because this reduces the effects of their errors through averaging. If different standard concentrations are used, doing so also enables the experimenter to test the instrument (or method) response linearity (e.g. see ref. [5]). However, this approach is limited by the constraint that each standard measurement costs time and materials. Often standard measurements replace the opportunity to measure unknown samples; for example, protein electrophoresis gels have a fixed number of lanes, so lanes that are used for standards cannot be used for unknown samples. Additionally, even though it is best to measure standards repeatedly, this doesn’t always happen in practice. Instead, for any of many possible reasons, data sets may contain valuable measurement data but insufficient standard measurements.

Calibration often needs to be performed repeatedly. For example, many experimental methods analyze samples in groups in which the sensitivity is the same for all measurements within a group but different for measurements in different groups (e.g. immunoblots and ELISA assays). Multiple calibrations are also required when one has many instruments that have different sensitivities. Additionally, most instrument sensitivities “drift” over time, necessitating periodic re-calibration (e.g. spectrometers and chromatographs). For convenience, we call all of these situations “batch-analyses,” defining a batch as any collection of measurements for which the sensitivities can be considered to be constant. By implication, each batch requires its own calibration.

We show here that calibrating each batch independently of the others, which is typical, is not the best approach, but that spreading sample replicates across different batches and then performing a simultaneous analysis of the data in all batches can substantially reduce the effects of measurement noise. In brief, our approach is to fit a statistical model to all of the data in a single step, finding both the instrument sensitivities and analyte amounts that best agree with all of the measurements. In other words, we cross-calibrate each batch against every other one. We call this the 1-step method, in contrast to the conventional 2-step method. The principle advantage of the 1-step method is that it makes calibration less sensitive to individual standard measurements. This often enables the use of batches that did not include any standards and it also enables the detection of errors in standard measurements. The results of the 1-step method are the maximum likelihood predictors, meaning that they are the results that are most probable within the assumptions of a statistical model.

We developed the 1-step calibration method to analyze data that we recently collected on proteins in mouse skin tumors. Our goal was to compare the relative levels of each of 7 different proteins (CypA, Hsp90, Hsp70, Hsc70, P53, Raf, and pERK) in 230 precancerous and cancerous mouse skin tumors using quantitative immunoblotting methods [6-10]. In brief, tumor extracts (replicated, pre-mixed with denaturing SDS-sample buffer, and stored at -80 C in small aliquots to maintain their integrity) were run on polyacrylamide gels (SDS-PAGE) to separate proteins by size and charge, followed by their transfer to nitrocellulose membranes. To individually probe query proteins of different molecular weights, the membranes were cut into horizontal strips bracketing size ranges determined by visible molecular weight standards that were run with each gel. Each strip, usually containing just one, or at most two query proteins of close molecular weight, was probed with the appropriate primary antibody (Spratt et al., in preparation). This was followed by incubation with secondary antibodies linked to an infrared fluorophore using the LICOR fluorescent Western blot detection system [11,12]. This method assured that signal intensity was linear within a large dynamic range (e.g. see [5]).

Calibrating these data was challenging for several reasons. First, immunoblotting is inherently imprecise. Indeed, all of the samples in our study, including those for standards, exhibited substantial measurement error (after calibration, our average CV was 37%). For this reason, we analyzed each sample multiple times on different blots so that we could reduce the effects of measurement noise through averaging. In total, we analyzed 230 tumor extracts on 117 immunoblots, each of which held up to 20 lanes (1510 replicated samples total, average of 6 replicates/sample). Secondly, one cannot directly compare fluorescence measurements between different blots because each blot’s sensitivity is strongly affected by minor experimental differences [9]. As a result, each blot needed to be treated as its own batch, with its own batch-specific sensitivity (calibration showed that they varied 27-fold between least and most sensitive). Finally, we could not use internal standards in this investigation (see [6]), which in this case would be naturally expressed proteins that are expected to have nearly constant concentrations such as the products of housekeeping genes, because tumors are very heterogeneous. As a result, we had to use a separate external standard, which was then subject to independent measurement errors. We created our standard by pooling several samples together to produce a single sample that included all of our proteins of interest [13].

Our 1-step calibration method is distinct from several other modifications to the classic calibration problem. Of particular note, Krutchkoff showed, nearly 50 years ago, that it can be better to fit the experimental results for the standard using the conversion function (eq. 2), rather than with the response function (eq. 1), which is called the inverse approach [14,15]. This led to an active debate about the relative merits of the two methods, along with the development of inverse regression methods [2,4,16]. From our reading of the literature, this debate appears to have largely ended by now, although without a clear winner. Other modifications to the classic calibration problem include Baysian [17] and non-parametric [3,18] methods. Bayesian methods are particularly helpful when the instrument is relatively insensitive to analyte variation (i.e. *β* is small) and the non-parametric methods when the measurement errors are substantially non-normally distributed. Finally, bootstrapping methods [19,20] can provide more accurate confidence intervals for the results, particularly for multivariate problems. In contrast to these developments, our 1-step approach follows the style of the classic calibration approach. It keeps the linear statistical model and the least squares fitting approaches, but simply extends them to account optimally for multiple batches. Our method builds on other analysis methods that also accounted for variability between batches [21-23] but, to the best of our knowledge, has not been described before. However, it is sufficiently straightforward that we would be surprised if some version of it has not been used previously.

## Results

### Definitions and model

Extending the analytical chemistry example given above, consider the situation in which one is quantifying the amount of an analyte in each of many samples, where a sample is simply some quantity of material. Assume this work is performed in batches, where a batch is a collection of measurements for which the instrument (or experimental method) sensitivity can be assumed to be constant. Additionally, assume that one or more standards are included in the analysis, where the standards already have well characterized analyte amounts. If such a standard is not available, then one simply assigns the role of the standard to one of the unknown samples and measures the other analyte amounts relative to that one. Our case followed this situation reasonably closely: the different mouse tissue extracts were our samples, the measured protein species in these samples were our analytes, the immunoblot gels were our batches, and the pooled sample served as our standard. This situation generalizes to many other calibration problems, too.

Assume that the following statistical model accurately represents the experimental data:

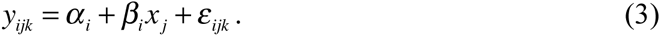

On the left side of the equation, each *y*_*ijk*_ value represents a single measurement, where *i* is the batch number, *j* is the sample number, and *k* distinguishes between multiple measurements of a particular sample that are within a single batch. Every measurement can be assigned a unique set of *i*, *j*, and *k* subscripts and so can be identified in this way. However, this does not necessarily imply that every sample was measured in every batch. To the contrary, most samples are likely to have been measured only a few times total in the entire experiment, making the *y*_*ijk*_ values a relatively sparse dataset (e.g. we had 230 total samples but only analyzed up to 20 at a time on any given immunoblot). On the right side of the equation, *α*_*i*_ and *β*_*i*_ are batch-specific sensitivity coefficients, *x*_*j*_ is the amount of analyte in sample *j*, and *ε*_*ijk*_ is the measurement error that arose in the *k*’th measurement of sample *j* in batch *i*. Assume that this error is normally distributed with mean of zero and standard deviation of *σ*, and that it is independent between measurements. This statistical model is very simple and builds upon conventional assumptions (including, importantly, that measurements depend linearly upon analyte amounts). It was also appropriate for our work because our immunoblot detection was linear in antigen amounts [12] and our tests of measurement repeatability showed reasonably independent and normally distributed errors (we found that the distribution of squared differences between repeated measurements of the same samples on the same blots was reasonably exponential, as one would anticipate for normally distributed errors). Table 1 summarizes the nomenclature introduced here.

**Table 1.** Data analysis nomenclature.

|  |  |
| --- | --- |
| **Roman symbols** |  |
| $a_i, b_i$ | estimates of batch sensitivity parameters |
| $i, N_B$ | batch index and number of batches |
| $j, N_S$ | sample index and number of samples |
| $k$ | measurement index within a specific batch and sample |
| $n_{i,j}$ | number of measurements of sample $j$ in batch $i$ (replacing $i$ or $j$ with “All” denotes all batches or samples) |
| $SD_j, SE_j$ | standard deviation and standard error for analyte amount $j$ |
| $T$ | list of standards |
| $x_j$ | analyte amount in sample $j$ |
| $y_{i,j,k}$ | measurement value for measurement $k$ of sample $j$ in batch $i$ |
| **Greek symbols** |  |
| $\alpha_i, \beta_i$ | true batch sensitivity parameters |
| $\varepsilon_{i,j,k}$ | measurement error for a specific measurement |
| $\sigma$ | standard deviation of measurement noise |
| $\chi^2$ | goodness of fit parameter |

The primary data analysis goal, typically, is to estimate the analyte amounts, *x*_*j*_, and their confidence intervals. Below, we also solve for the sensitivity coefficients, *a*_*i*_ and *b*_*i*_, which can enable one to calibrate any new measurements that were not included in the original data. We also find the measurement standard deviation, *σ*, which can be helpful for improving the measurement technique and for identifying any outlier data points.

### The 2-step method

We present the conventional 2-step calibration method, focusing on its application to samples that are measured in batches, to introduce our mathematical notation in a setting that may be familiar and to show some aspects of the method that are widely overlooked. The left side of Figure 1 illustrates the 2-step method.

**Figure 1.**
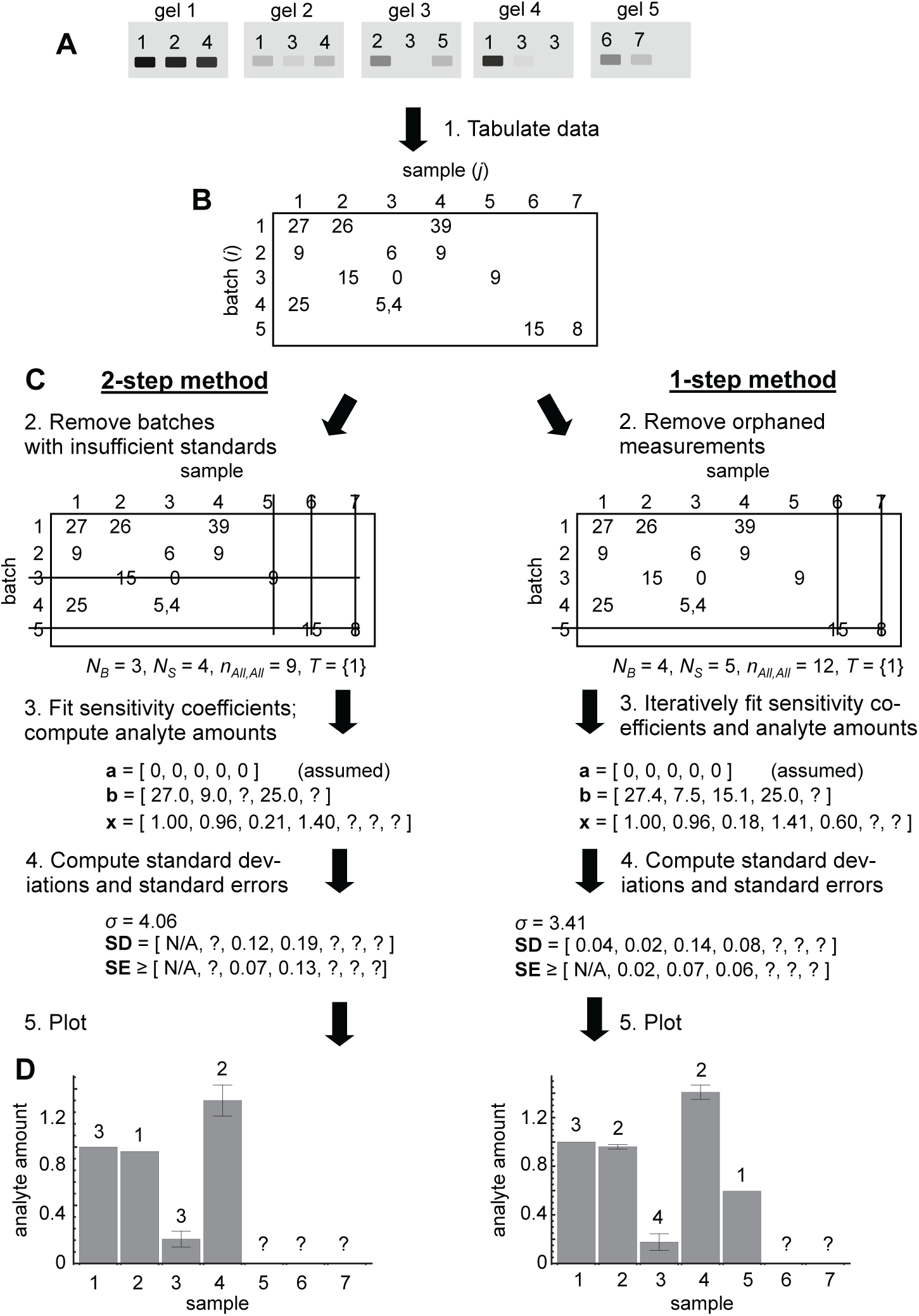
Comparison of workflow for 2-step and 1-step calibration methods, illustrated for calibrating band intensities on immunoblots. A. Illustration of samples 1 (the standard) through 7, run on 5 different immunoblots with variable replication. The band intensities shown depend on the sample, blot, and experimental noise. B. Tabulated data showing assigned band intensities for each sample and blot. C. Direct comparison of the conventional 2-step calibration method (left) with the 1-step calibration method (right). D. Plots of the calibrated estimates of analyte amounts in each sample using the different methods. Error bars represent the standard error of the mean and numbers above the bars represent the number of calibrated measurements of each sample.

#### Tabulate data

The measurements need to be tabulated, putting each sample in a separate column and each batch in a separate row. Each table site has as many entries as there are measurements for that specific sample and batch, which may be zero, one, or more than one.

#### Remove batches with insufficient standards

To enable calibration, each batch needs to include at least as many different standard measurements as there are unknown sensitivity coefficients (because of the linear algebra result that one needs at least *n* equations to solve for *n* unknowns). The statistical model (eq. 3) includes two sensitivity coefficients, *α*_*i*_ and *β*_*i*_, so each batch generally needs to include at least two different standard measurements. On the other hand, if one assumes that measurements do not have a consistent offset, meaning that all of the *α*_*i*_ values are assumed to equal zero, then each batch only needs one standard measurement. Figure 1 illustrates this latter situation. Our work also fit this latter situation because we corrected for background fluorescence before starting our data calibration. Any batches that do not include as many standard measurements as unknown sensitivity coefficients need to be removed from the data analysis. In the process, any samples that were only measured in these batches get removed too.

Next, it is helpful to define several variables. Define *N*_*B*_ as the number of batches (number of rows), *N*_*S*_ as the number of samples (number of columns), and *n*_*ij*_ as the number of measurements of sample *j* in batch *i* (number of entries at site *i,j*).

Generalizing this last definition, *n*_*All,j*_ is the total number of measurements of sample *j* (the number of entries in column *j*), *n*_*i,All*_ is the total number of measurements in batch *i* (the number of entries in row *i*), and *n*_*All,All*_ is the total number of measurements (the number of entries in the table). Also, define *T* as the list of standards; for example, there is one standard in Figure 1, which is sample number 1, so *T* = {1} in that case. Finally, *n*_*i,T*_ is the number of standard measurements in batch *i*.

#### Fit sensitivity coefficients

As the first step of the 2-step method (the calibration stage), a line is fit to the standard data in each batch using least-squares methods. This provides best-fit *a*_*i*_ and *b*_*i*_ values as estimates for the “true” *α*_*i*_ and *β*_*i*_ sensitivity coefficients. If the *α*_*i*_ sensitivities are not assumed to equal zero, then the *a*_*i*_ and *b*_*i*_ values are found using the standard results for simple linear regression [24],

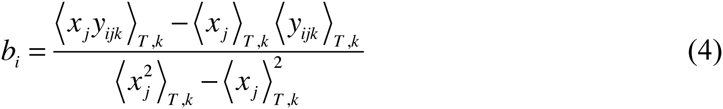

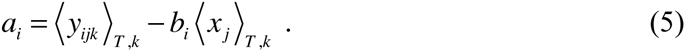

Angle brackets indicate averaging over the indices that are listed in their subscripts. In this case, the average is over all standards that were measured in any particular batch. For example,

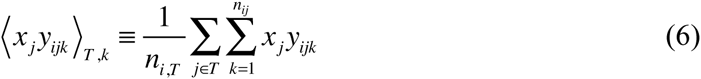

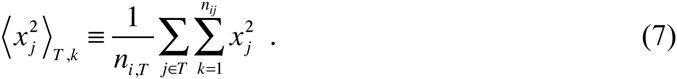

If the *α*_*i*_ sensitivities are assumed to equal zero, then all of the *a*_*i*_ values clearly equal zero and the *b*_*i*_ values simplify to

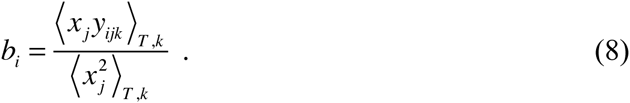

Note that an intuitively sensible, but incorrect, approach would be to compute the *b*_*i*_ values in the latter case by simply solving *y*_*ijk*_ ≈ *b*_*i*_*x*_*j*_ for *b*_*i*_ to give *b*_*i*_ ≈ *y*_*ijk*_/*x*_*j*_ and then averaging these values to give *b*_*i*_ = <*y*_*ijk*_/*x*_*j*_>_*T,k*_. Eq. 8 is different in that it weights each term in this average by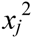. Doing so correctly emphasizes those data points that are likely to have larger measurement values and hence lower relative errors (see the derivations in the appendix).

##### Compute analyte amounts

In the second step of the 2-step method (the prediction stage), the amount of analyte in each unknown sample is computed by inverting the statistical model equation (eq. 3), while using the *a*_*i*_ and *b*_*i*_ estimates for *α*_*i*_ and *β*_*i*_. Then, averaging results over all analyses of each sample yields the following estimate for the sample’s analyte amount:

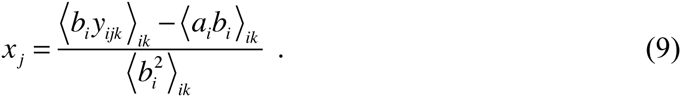

As in eq. 8, this solution is weighted to emphasize the data points that have larger measurement values and hence lower relative errors. In contrast, the intuitively sensible but incorrect approach gives the average as *x*_*j*_ = <(*y*_*ijk*_–*a*_*i*_)/*b*_*i*_>_*ik*_, but this over-emphasizes data points that are likely to have large errors and under-emphasizes those that are likely to have small errors.

#### (4) Compute standard deviations and standard errors

Our statistical model assumes that measurements have normally distributed errors. To estimate the standard deviation of those errors, we compute the root mean square (rms) average deviation of the actual measurements, *y*_*ijk*_, away from where we would have expected them, *a*_*i*_+*b*_*i*_*x*_*j*_,

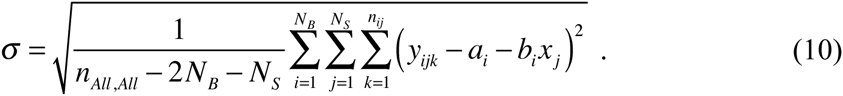

The denominator represents the number of degrees of freedom, which is one for each of the *n*_*All,All*_ data points, minus the number of fit coefficients. There are 2*N*_*B*_+*N*_*S*_ fit coefficients if the *α*_*i*_ values are not assumed to equal zero (for the *a*_*i*_, *b*_*i*_, and *x*_*j*_ values), as shown in eq. 10, and *N*_*B*_+*N*_*S*_ if the *α*_*i*_ values are assumed to equal 0. Because we assumed Gaussian distributed noise, about 68% of the measurements should be within one standard deviation of their expected values and about 95% within two standard deviations. Measurements that are many standard deviations away from their expected values are outliers, which may warrant further inspection and possible removal.

Importantly though, if the minimum number of standards were measured in each batch, which is typical, then it is impossible to determine if any of them are outliers because the sensitivity parameters were computed directly from their measurements.

Separate standard deviations represent the variability in the different analyte amount estimates, which came from eq. 9. These estimates are weighted means, so their variabilities are computed as weighted standard deviations, for which the general equation is [25]

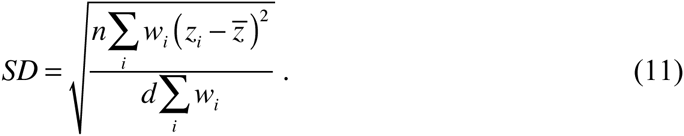

Here, *z*_*i*_ represent the data, *w*_*i*_ represent the weights, *z* is the sample mean, *n* is the number of data points, and *d* is the number of degrees of freedom. Applying this to the sample analyte amounts and simplifying gives

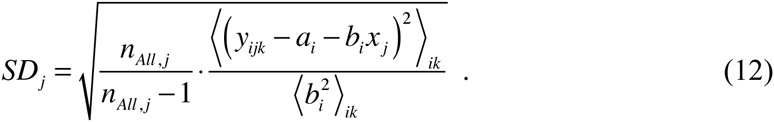

The number of degrees of freedom is *n*_*All,j*_-1 because there are *n*_*All,j*_ terms in the sum but the *x*_*j*_ value was constrained through eq. 9.

The standard errors of the means reflect the accuracy with which the *x*_*j*_ values are likely to represent the true analyte amounts. As usual, they are computed by dividing the standard deviations by the square root of the number of measurements being considered [25]. However, doing so yields a lower bound for the standard error because the standard deviations were computed while assuming that the *a*_*i*_ and *b*_*i*_ values equaled their true values and that the *x*_*j*_ value was the only one that needed to be fit to the data. However, all three of these are estimates, which increases the uncertainty for the analyte amounts. Thus, the standard errors are

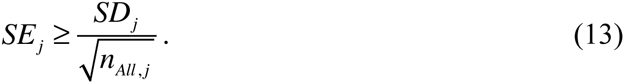

The interpretation is that the difference between each computed *x*_*j*_ value and the true analyte amount for the sample is likely to be a Gaussian distributed random variable with standard deviation equal to *SE*_*j*_. This result does not apply to the standards because their analyte amounts are assumed to be known.

## 1-step method

The 1-step method parallels the 2-step method very closely.

1. *Tabulate data.* The 1-step method uses the same data table as the 2-step method.
2. *Remove orphan measurements.* The 1-step method relies on standards less than the 2-step method does, but still requires that each measurement can be related to the standard measurements in some way. More precisely, each batch needs at least as many independent “connections” to standard measurements as there are sensitivity coefficients; a batch is connected to a standard if (*i*) it includes a measurement of that standard or (*ii*) it shares a sample with some other batch that is connected to that standard. We call measurements that cannot be connected to enough standard measurements orphans. These orphan measurements need to be removed from the data analysis, along with the samples and batches to which they belong. The 1-step method uses the same definitions for the *N*_*B*_, *N*_*S*_, *n*_*i,j*_, *T*, and other variables as the 2-step method.
3. *Iteratively fit sensitivities and analyte amounts.* The single step of the 1-step method is to simultaneously fit the *a*_*i*_, *b*_*i*_, and *x*_*j*_ values to the data while assuming the statistical model given in eq. 3. This can be accomplished in many ways, including with deterministic and stochastic minimization algorithms [24]. However, we found that computing the sensitivities and analyte amounts iteratively, using equations derived in the appendix, was particularly simple and efficient. In this method, one first guesses all of the sensitivities. An adequate approach is simply to set all of them to 1 initially, but we found that results converged faster when we guessed as many as possible using eqs. 4, 5, and 8 from the 2-step method and then set the rest to their means. Next, the unknown analyte amounts are computed from

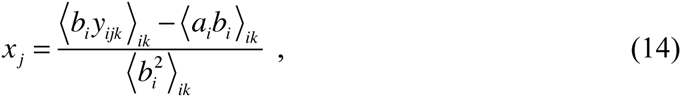

which is identical to eq. 9. Then, the sensitivities are computed from

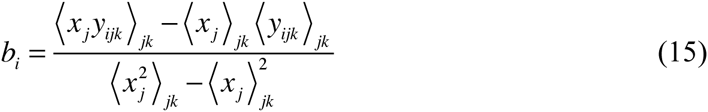

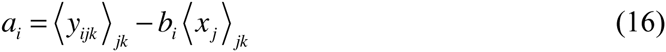

if the *α*_*i*_ values are not assumed to equal zero, and

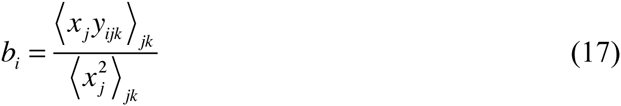

if they are. These equations only differ from eqs. 4, 5, and 8 in that they include averages over all measurements in a batch rather than just the standard measurements. Iterating over eqs. 14 to 17 leads to the best-fit values for the analyte amounts and sensitivities. We continued until all sensitivity parameter and analyte amount estimates changed by less than 1 part in 10^5^ between subsequent iterations, which never took more than a few hundred iterations (340 for our immunoblot data and about 70 for most of the validation tests described below).
4. *Compute standard deviations and standard errors.* The standard deviation and standard error equations that are presented above in eqs. 10 to 13 apply here as well. However, the standard deviation *can* be used to identify outlier standard measurements in this case, even if relatively few standard measurements were made, because these sensitivity parameters were computed from all of the measurements instead of just the standard measurements.

## Validation

We validated our method by analyzing 1000 artificially generated data sets and comparing the fit results with the true parameters from which the data sets were generated. Each data set comprised 20 samples that had random analyte amounts, 20 batches that had random *α* and *β* sensitivity values, and 400 measurements that were randomly distributed in the data table. The first two samples were assigned to be standards, with fixed analyte amounts. We analyzed each data set with both the 1-step and 2-step methods. Figure 2A-D shows results from one of these data sets. It shows that both analysis methods were able to recover the true parameters from the data reasonably well, but the 1-step method generally led to smaller errors. There were enough standard measurements in this data set that all analyte amounts could be estimated using both methods. However, only 8 of the batch sensitivities could be estimated using the 2-step method because the others had insufficient standards and so were removed from the analysis (note the relatively few gray data points in panels C and D).

**Figure 2.**
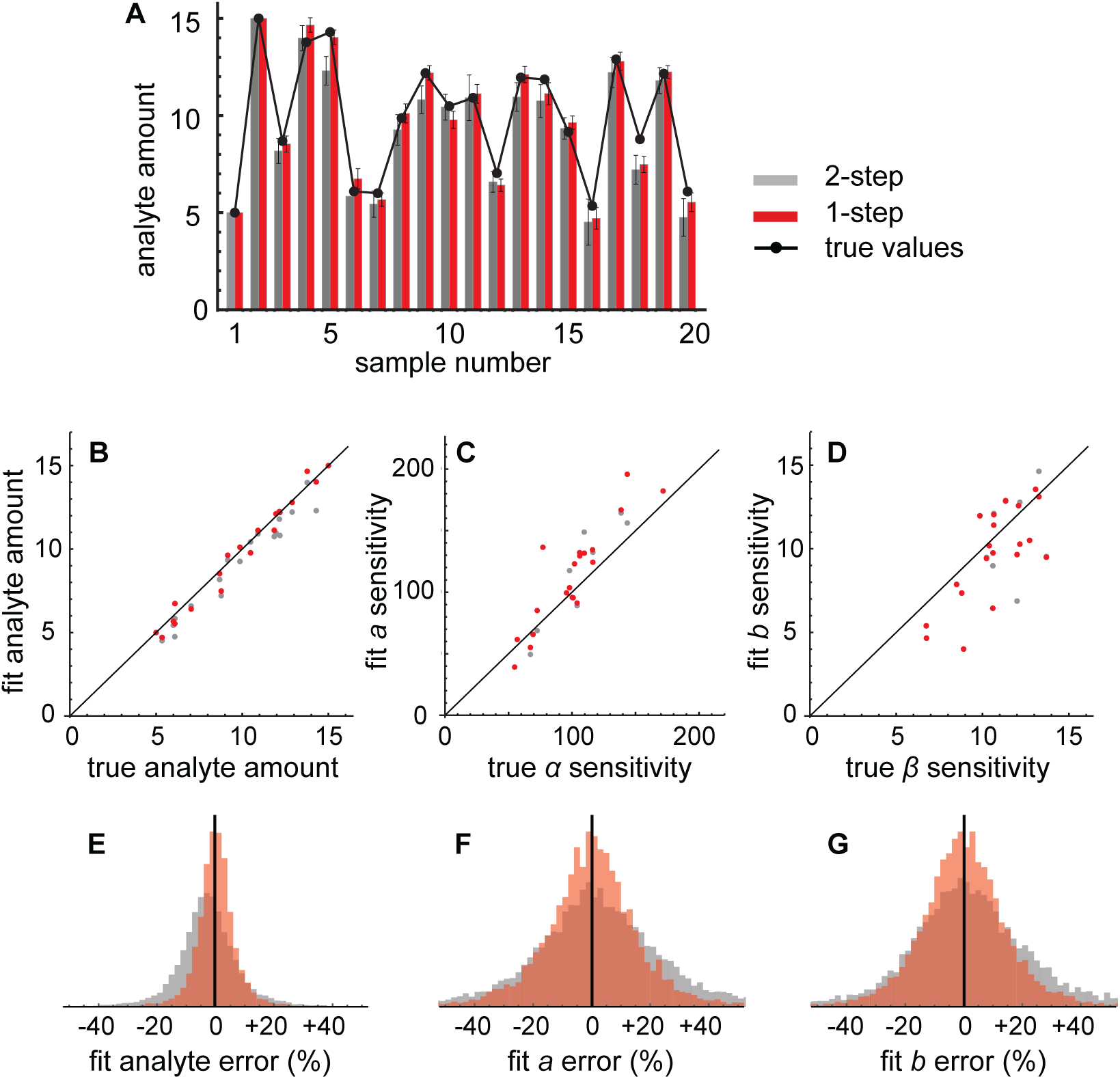
Comparison of the 1-step and 2-step methods using artificial data sets. A. Sample analyte amounts for an artificial data set; see the main text for details. Here and in subsequent panels, black features represent the true analyte amounts, gray features represent results from the 2-step method, and red features represent results from the 1-step method. Error bars represent standard errors. (B-D) Comparison of computed sample analyte amounts, *a* sensitivity coefficients, and *b* sensitivity coefficients with their true values for the same artificial data set. (E-G) Histograms of errors between fit values and true values for computed sample analyte amounts, *a* sensitivity coefficients, and *b* sensitivity coefficients for 1000 artificial data sets. In all cases, the 1-step method yields more accurate data calibration. Data sets were generated with: 20 batches with Gaussian distributed *α* values with mean of 100 and standard deviation of 30, 20 samples with Gaussian distributed *x* and *β* values with mean of 10 and standard deviation of 3, 400 measurements distributed randomly in the data table with *σ* set equal to 20, and standard analyte amounts set to 5 and 15.

We observed essentially the same results for the other artificial data sets as well. Figure 2E shows that the 1-step method overestimated analyte amounts by 0.2% on average, whereas the 2-step method underestimated them by 2.6% on average. Further tests showed that these deviations arose from the choices of standards, becoming larger when the standard analyte amounts differed more from typical sample analyte amounts. However, the 1-step method always had much smaller deviations. Figure 2E also shows that the 1-step method generally computed individual analyte amounts that were closer to the true values: the root mean square (rms) error for the analyte amounts was 8% for the 1-step method and 13% for the 2-step method. Similarly, Figure 2F-G show that the 1-step method computed sensitivity parameters that were closer to their true values: rms errors were 18% and 27% for the *a* sensitivity parameter and 17% and 25% for the *b* sensitivity parameter, for the two methods respectively (this comparison only includes parameters that both methods computed successfully, to make them comparable).

Analysis of the results showed that these improvements arose from two factors. First, the 1-step method included more data points in the calibration due to its decreased dependence on standards (out of the 1000 data sets, none of the batches needed to be removed from the analysis in the 1-step method but 60% of them needed to be removed for the 2-step method). As a result, the 1-step method was able to include more measurements in its averages and hence reduce the effects of measurement noise. Secondly, the 1-step computed the sensitivity parameters more accurately, even when there were sufficient standards for both methods. To investigate this latter point further, we repeated the validation procedure but altered it so that every batch included every standard. As a result, no measurements needed to be removed during either analysis. In this case, rms errors for the analyte amounts were 14% and 16% for the 1-step and 2-step methods, respectively, again showing smaller errors for the 1-step method.

We also compared the computed standard deviations and standard errors against the true ones as a way of validating eqs. 9 and 13. We found good agreement. The 1-step and 2-step methods estimated the measurement standard deviation to be 17 and 21 units, respectively, while the true value was 20 units. Also, the average 1-step and 2-step standard error estimates were 76% and 71% of the actual deviations between the computed and true analyte amounts. These show reasonable agreement and are consistent with the inequality in eq. 13.

## Protein immunoblot data

We analyzed our experimental immunoblot data using both methods, of which a small portion of the results are shown in Figure 3. These data are scaled so that the standard (not shown in the figure) has an analyte amount of 1. As part of the analysis, we automatically removed all measurement results that were 4 or more standard deviations away from their expected values, which we deemed to be outliers, and then re-calibrated the remaining data until there were no more outliers. This process showed that about 1% of our measurement results were outliers (for comparison, 0.003% would be expected to be more than 4 standard deviations away from the mean if errors were distributed perfectly normally). After all outliers were removed, the 1-step method enabled us to use all of our measurements in the final analysis, whereas we would have needed to remove about 35% of them with the 2-step method.

**Figure 3.**
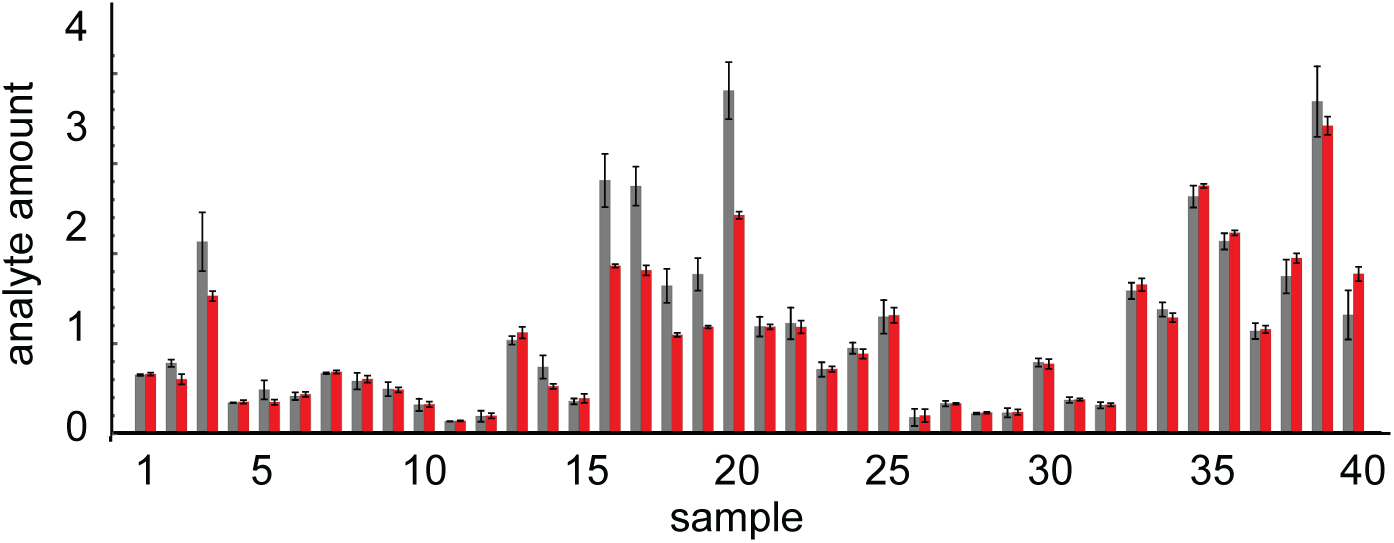
Calibrated experimental immunoblot data. This figure shows calibrated analyte amounts for 40 of our 230 samples that we analyzed with immunoblots. The others were qualitatively similar. Gray bars represent results from the 2-step method, red bars represent results from the 1-step method, and error bars represent standard error values. On average, there were 3.4 calibrated measurements for each sample with the 2-step method and 4.9 for the 1-step method. Note that the 1-step method results have smaller standard errors.

Differences between the two methods were more striking with the real data than with the artificial data that we used for validation. Here, the two methods often returned substantially different analyte amount estimates. Also, the 1-step method typically returned substantially smaller standard errors for the analyte amount estimates, with a mean standard error of 20% as compared to 31% for the 2-step method. We are using the 1-step method results for further investigation of these data.

## Discussion

We have described a method for calibrating data to external standards. The conventional approach to calibrating measurement data, which we call the 2-step method, is justifiably nearly ubiquitous. It is simple, intuitive, and convenient. As a result, it can be performed by hand or with spreadsheet software. Also, if there is only a single batch of data, then it is the optimal approach, returning the maximum likelihood predictors for the analyte amounts (assuming that the statistical model is correct and that the measurements are weighted properly when averaging, as shown above). However, it does not return the best possible results if there are multiple batches because it ignores information from samples that were measured in multiple batches. This makes it particularly sensitive to errors in the standard measurements, and also completely reliant on there being sufficient standards in every batch. On the other hand, our novel 1-step method uses information from samples that were measured in multiple batches. This decreases its reliance on standard samples and enables it to return more accurate results, which are the maximum likelihood predictors, now for the complete data set.

A drawback of the 1-step method is that it requires an iterative computation, making it impractical to perform by hand or in a simple spreadsheet. Nevertheless, this computation is not particularly demanding. Calibrating our immunoblot data set, which comprises 5966 measurements and requires 340 iterations, takes just over 1 minute on a 2013 MacBook laptop computer. From inspection of eqs. 14 to 17, the computational demands scale approximately linearly with the number of measurements, implying that much larger data sets can be calibrated reasonably efficiently as well. A second drawback of the method is that it assumes that instrument or method responses increase linearly with analyte amounts (see eq. 3), which is often not the case. However, it is relatively straightforward to modify the 1-step method as it is presented here to specific non-linear relationships by repeating the derivations presented in the appendix, but for the desired relationship.

An obvious question arises of how to best design experiments so that they yield the most accurate results, while calibrating the data with the 1-step method. Although addressing it was beyond the scope of our work, some aspects are reasonably obvious.

First, we anticipate that it is best to measure standards in as many of the batches as possible because that minimizes the number of steps that need to be taken to connect unknown sample measurements with standard measurements. Also, we suspect that it is better to spread replicates of sample measurements out over multiple batches, rather than to perform them all within a single batch, because that improves the ability to cross-calibrate the different batches.

Our software for calibrating data using both the 1-step and 2-step methods is written in Python, is open source, and is in the public domain (i.e. we do not reserve any intellectual property rights). It is available at http://www.smoldyn.org/calibration.html. It can also be used at the same website as an online calibration service.

## Appendix

This appendix derives most of the equations presented above. It is shown at a relatively elementary level to make it widely accessible, so statistics textbooks (e.g. ref. [25]) should be consulted for more rigorous treatments.

From eq. 3, we assume the statistical model

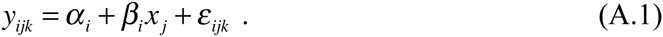

We rearrange the equation and divide both sides by *σ*, the measurement error standard deviation, to yield the scaled measurement errors,

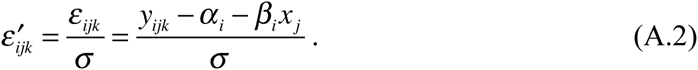

Because we assumed that the measurement noise is Gaussian distributed and independent between data points, the 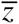 values are independent normally distributed random variables with zero mean and unit standard deviation. We square both sides of this equation and sum over all data points to yield

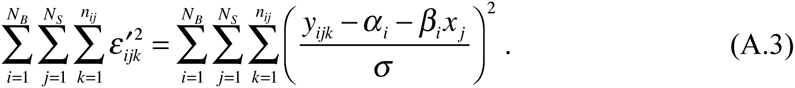

The left side is a sum of squared independent normally distributed random variables, which means that it is itself a random variable and it obeys the chi-squared distribution.

Looking back at eq. A.1, if we knew the exact values of each *α*_*i*_, *β*_*i*_, and *x*_*j*_ but not the *y*_*ijk*_ values, then the assumption that the error is normally distributed with a mean value of zero would imply that the most likely value for *y*_*ijk*_ is the one that arises if the error equals zero. However, we actually know the *y*_*ijk*_ values but not the *α*_*i*_, *β*_*i*_, or *x*_*j*_ values. So, we rearrange the prior statement to claim that the most likely values of *α*_*i*_, *β*_*i*_, and *x*_*j*_, given the known *y*_*ijk*_ values, are those that minimize the computed errors (eq. A.2). This rearrangement is not completely legitimate but is the central ansatz of maximum likelihood estimation and is partially justified by Bayesian analysis [24]. Without going further into the details, we perform maximum likelihood estimation by replacing the true sensitivity coefficients, *α*_*i*_ and *β*_*i*_, in eq. A.3 with the unknown *a*_*i*_ and *b*_*i*_ estimated sensitivity coefficients to yield the following “goodness-of-fit” function,

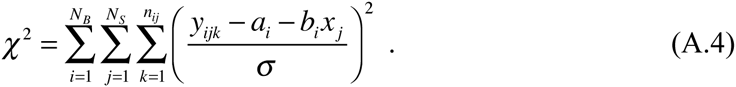

We then minimize this function with respect to each *a*_*i*_, *b*_*i*_, and unknown *x*_*j*_ parameter to find their most likely values. The parameter values that minimize the *χ*2 function are called the maximum likelihood predictors because they are the most probable values, within the assumptions of the model.

We find the minimum of *χ*2 with respect to *x*_*j′*_, where *j′* is the index of a specific unknown sample, by differentiating *χ*2 with respect to *x*_*j′*_ and setting the result to zero:

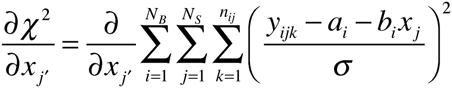

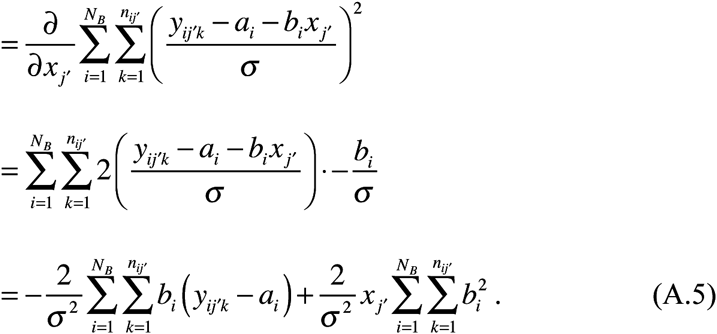

Setting the result to zero, renaming *j′* to *j*, and simplifying yields

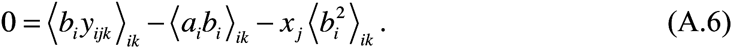

This result represents one equation for each unknown sample. Minimizing *χ*2 with respect to *a*_*i*_ and *b*_*i*_ are analogous, yielding

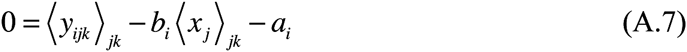

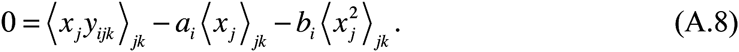

These results represent one pair of equations for each batch. In principle, equations A.6 to A.8 can be solved for the unknown *a*_*i*_, and *b*_*i*_, and *x*_*j*_ values. However, this appears to be analytically intractable so instead we rearrange them to yield

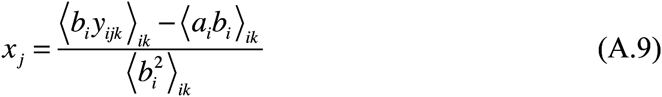

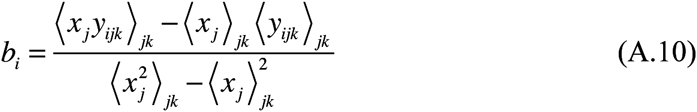

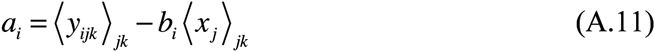

If it is assumed that the *α*_*i*_ values all equal zero, then the *a*_*i*_ values are set to zero and the solutions for *x*_*j*_ and *b*_*i*_ get simplified to

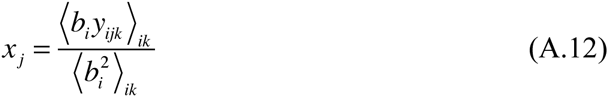

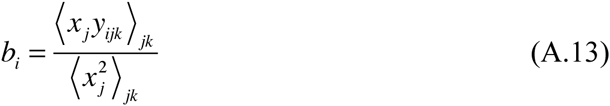

These equations cannot be computed sequentially because each equation requires knowledge of the other results. Thus, the approach taken by the 2-step method is to limit the averages in eqs. A.10, A.11, and A.13 to just those samples which have known analyte amounts, which are the standards. After this, eqs. A.9 or A.12 can be computed without problems. Alternatively, the approach taken by the 1-step method is to compute the equations iteratively, which then yields the best-fit *a*_*i*_, *b*_*i*_ and *x*_*j*_ values. To convince ourselves that the iterative method leads to the correct solutions, we also minimized *χ*2 using Mathematica’s “NMinimize” function for a series of validation data sets. In all cases, results were identical but the iterative approach was many-fold faster.

To compute the measurement standard deviation, we start with the fact that the mean of a chi-squared distribution is equal to the number of random variables that are summed. In eq. A.4, the *χ*2 sum includes *n*_*All,All*_ terms, suggesting that this would be the mean of the distribution. However, we don’t know the true *α*_*i*_, *β*_*i*_, or *x*_*j*_ values, but only those that we fit by minimizing *χ*2, which reduces the mean by 2*N*_*B*_+*N*_*S*_ degrees of freedom. Using the assumption that any specific data set is likely to be reasonably typical, we equate *χ*2 to *n*_*All,All*_–2*N*_*B*_–*N*_*S*_, yielding

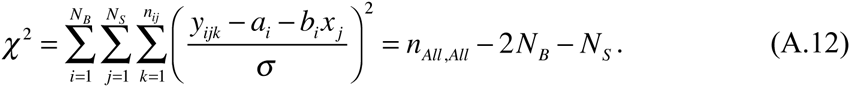

Solving for the measurement standard deviation then yields

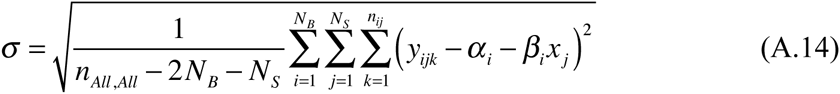

Finally, we solve for the individual sensitivity coefficient and analyte standard deviations. Both are simply weighted averages, so we use the general equations for a weighted standard deviation (main text eq. 11) to yield the results

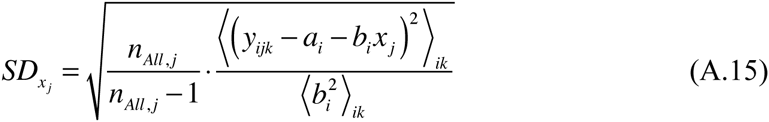

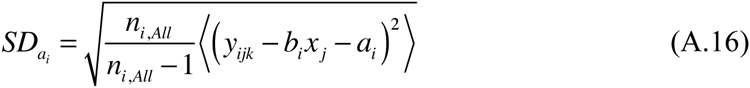

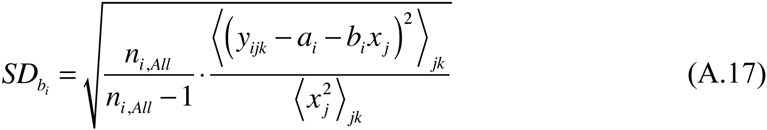

Dividing these results by the square root of the number of data points yields estimates for the standard errors.

## Acknowledgements

We thank Roger Brent for valuable discussions and critical comments on a previous version of the manuscript, along with funding that he provided for SSA. We also thank Wenying Shou and Sam Oman for helpful discussions. We are grateful for funding from several sources: NIGMS grants R01GM086615, awarded to Roger Brent and Richard Yu, and R01GM097479, awarded to Roger Brent, funded SSA; also a Scholar Award of the Damon Runyon-Walter Winchell Foundation and NIGMS grant R01GM068873 funded SR.

